# Loss of kinase Atg1 increases yeast maintenance energy requirement

**DOI:** 10.1101/2023.03.06.531285

**Authors:** Xin Chen, Aafke C. A. van Aalst, Dina Petranovic, Markus M.M. Bisschops

## Abstract

Maintenance of cellular homeostasis underlies healthy aging. The processes involved in homeostasis rely on the so-called maintenance energy requirement and changes in this maintenance energy requirement impact aging and survival. Among maintenance processes, autophagy plays a crucial role as it is involved in the turn-over and recycling of damaged cellular material, such as organelles or proteins. The contribution of autophagy to the maintenance energy requirement is however unknown. Taking advantage of the high degree of conservation of autophagy between humans and *Saccharomyces cerevisiae*, we have used this yeast as a model organism to study the impact of macroautophagy on the maintenance energy requirement. The combination of the GFP-Atg8 cleavage assay with yeast retentostat cultures showed that autophagy is highly active in chronologically aging yeast cells, in non-dividing, but non-starving conditions. Deletion of the autophagy-activating kinase *ATG1*, homolog of human *ULK1*, resulted in a 60% increase in the maintenance energy requirement and doubled the specific death rate. Both these increases cannot be solely attributed to an observed increase in loss of respiratory capacity. Intriguingly, loss of Atg1 only reduced GFP-Atg8 cleavage by 20% under these conditions, indicating that Atg1-indendent modes of autophagy might be active. Overall, we illustrate the importance of autophagy on the energetics of aging cells and propose an alternative system for the widely applied yeast stationary phase cultures in chronological aging studies.

## Introduction

Cellular maintenance and cellular aging both rely on processes that clear or repair damaged cellular components. Although their origins are debated, such damaged macromolecules and organelles are a hallmark of aged cells that enter senescence or face cell death (Barja 2019). Autophagy is one of the main routes for cells to turnover damaged macromolecules, e.g., proteins, organelles *etc.*, and therefore plays a crucial role in maintenance of cellular homeostasis. In addition to turnover, autophagy also leads to recycling of building blocks of these cell parts, such as amino acids, for biosynthetic purposes (Wen & Klionsky 2016). Macroautophagy is a type of autophagy which involves the formation of autophagosomes around cytosolic cargo, which is subsequently delivered to lysosomes/vacuole for degradation (Wen & Klionsky 2016). Macroautophagy is the most studied process of autophagy, and we will refer to macroautophagy and autophagy interchangeably in this study.

The yeast *Saccharomyces cerevisiae* is a unicellular model organism widely used to study different eukaryal processes, including cellular aging (Longo et al. 2012; Mirisola et al. 2014). Many processes linked to cellular maintenance and aging are conserved from yeast to human, for example signaling cascades such as the TOR pathway (Virgilio & Loewith 2006). This yeast model also strongly contributed to our current understanding of autophagy as this process appears to be highly conserved at morphological and mechanistic level (Wen & Klionsky 2016). This includes the only known kinase, Atg1 (Ulk1/2 in mammals), which is involved in autophagosome formation (Mizushima et al. 2011). Atg1 plays a vital role in autophagy (in complex with Atg13 and Atg17) as well as in the biosynthetic Cvt pathway (Kabeya et al. 2005; Takeshige et al. 1992). Maintenance of cellular homeostasis is also at the core of healthy aging. In this light, it may not be surprising that different interventions that enhance autophagy result in increased longevity, for example inhibition of the TOR complex and nutrient restriction (Tyler & Johnson 2018). The inverse also holds: inhibition of autophagy reduces healthy lifespan (Sampaio-Marques et al. 2014). This requires a fine balance between activation and modulation of autophagy activity and it is subject to complex and tight regulation (Delorme-Axford & Klionsky 2018).

To ensure their integrity, functionality and survival, cells have to invest energy in the so-called endogenous metabolism or maintenance energy requirement (Pirt 1982; Bodegom 2007). The maintenance energy requirement (*m*) for these processes highly dependents on the cell type and the conditions cells experience and can be expressed in energy equivalents (for example ATP; *m_ATP_* or glucose; *m_S_*) consumed per cell per unit of time (Bodegom 2007). Several core processes and mechanisms have been identified as part of maintenance, for example maintenance of proteostasis and ion-gradients, including proton motive forces (Sampaio-Marques & Ludovico 2018; Bodegom 2007). However, about the individual contributions of those processes to the maintenance requirement is much less known. One of the reasons for this is that under most conditions, cellular maintenance only makes up a small fraction, especially flux-wise, of the total fluxes within a cell (Ercan et al. 2015). In yeast, as in most microorganisms, the maintenance requirement is generally overshadowed by cell growth. Estimations of the glucose consumption for maintenance on a defined medium (0.04-0.5 mmol/g_cells_/h) (Vos et al. 2016; Boender et al. 2009) correspond to less than 5% of the total glucose consumption during fast exponential growth (13 mmol/g_cells_/h) (Chen et al. 2017). Although this energy consumption by maintenance processes seems small, it is of vital importance.

*S. cerevisiae* is used to study both replicative and chronological aging (Longo et al. 2012). Replicative aging and the concomitant replicative life span (RLS), the number of divisions a cell can undergo, is generally studied under conditions of fast growth. On the contrary, chronological aging and a cell’s chronological life span (CLS) are primarily studied in non-dividing cells (Mirisola et al. 2014). The current paradigm of non-growing and non-dividing yeast cells are stationary phase (SP) cultures (Gulli et al. 2019). In these cultures, growth arrest is caused by the depletion of one or more essential nutrients from the medium, often the carbon- and energy-source, such as glucose (Werner-Washburne et al. 1996). These starvation conditions enforce cells to use their intracellular energy reserves, such as glycogen and lipids, to fulfil the maintenance energy requirement. These reserves are however finite and consequently the timespan that maintenance processes can be fueled is limited. This can significantly interfere when trying to control conditions to study aging regulation and processes (Bisschops et al. 2015). Quantification of maintenance energy consumption rates and contributions of individual maintenance processes under starvation conditions is also challenging as it faces the difficulty to absolutely quantify intracellular metabolite concentrations. In addition, SP cultures are heterogeneous: cells differ in terms of survival and metabolic state (i.e., respiration *versus* fermentation), which has to be taken into account when interpreting data (Allen et al. 2006).

To overcome these drawbacks of classical SP cultures, retentostat cultures can be used. In a retentostat, cells are retained and accumulate inside a bioreactor with a continuous and precise energy (glucose) supply (Vos et al. 2016; Boender et al. 2009; Bisschops et al. 2017). The increasing cell number causes the continuously supplied glucose to be gradually divided over more and more cells. Ultimately the amount of glucose available per cell per time (e.g. g_glucose_/g_cells_/h), reduces to the maintenance requirement and no glucose is available for divisions anymore (Ercan et al. 2015). As a result, cell divisions cease in the population and in a typical yeast retentostat culture the specific growth rate decreases from 0.025 h^-1^ to below 0.001 h^-1^, corresponding to an increase in average cell division time from one day to over one month (Boender et al. 2009; Vos et al. 2016). Retentostat cultures also allow to accurately determine maintenance energy requirements as has been done for *S. cerevisiae* under aerobic and anaerobic conditions, as well as for other fungi and bacteria (Vos et al. 2016; Boender et al. 2009; Ercan et al. 2015). The conditions, which are *de facto* prolonged extreme calorie restriction (ECR), furthermore result in a higher cellular activity and longer survival compared to SP cultures. These characteristics make yeast retentostat cultures a promising tool to study chronological aging (Bisschops et al. 2017).

Previous genome wide-expression analyses of yeast retentostat cultures identified large sets of genes whose expression changed significantly. Notably, levels of 768 and 980 transcripts increased, while growth came to a virtual standstill, under aerobic or anaerobic conditions, respectively (Vos et al. 2016; Boender, van Maris, et al. 2011). Among these transcripts, several encode for autophagy-related proteins, involved in for example the general autophagy machinery (e.g., *ATG1*, *ATG10*, *ATG5*, *ATG7* and *ATG9*) or selective autophagy (e.g., *ATG11*, *ATG34*, *ATG36* and *ATG39*) (Farré & Subramani 2016). Transcript levels of *ATG22*, for example, increased 2.5-fold under increasing calorie restriction. *ATG22* encodes a vacuolar membrane protein involved in amino acid efflux and thereby recycling (Yang et al. 2006). Based on these observations, autophagy was hypothesized to be active and putatively play an important role in carbon-limited retentostat cultures, even though these cultures are not starved and actually show clear differences from their starved counterparts (Bisschops et al. 2015; Bisschops et al. 2017; Boender, Almering, et al. 2011).

In this study, we explore the role of autophagy in these non-dividing cultures. We selected this maintenance process as it is strongly linked to both replicative and chronological aging (Tyler & Johnson 2018). In addition, autophagy is expected to directly affect maintenance energy requirements: It may save energy by recycling building blocks for the cells or increase energy costs due to less/non-specific degradation of cellular components.

## Results

### Autophagy activity stays high in aging retentostat cultures

To monitor autophagy activity in different yeast cultures, the GFP-Atg8 cleavage assay was used (Cheong & Klionsky 2008). Atg8 is a component of autophagosomal membranes and is degraded upon delivery to the vacuole. GFP is more stable and a higher ratio of free GFP over total GFP (GFP-Atg8 plus free GFP), indicates stronger cleavage and degradation of the Atg8 moiety corresponding to active autophagy. The *gfp-ATG8* expression cassette under the control of the *ATG8* promoter was integrated in the genome of a prototrophic *S. cerevisiae* strain. Integration was chosen to avoid interference of auxotrophies or plasmid copy number variation in the assay and the molecular responses of cells. To the best of our knowledge, free GFP/total GFP ratios have not been determined yet in continuously fed yeast cultures. To be able to compare results from such cultures, the resulting strain IMI442 was first assayed under conditions of known low or high autophagy activity. During fast growth on both rich (YPD) and defined medium, low free GFP/total GFP ratios of 0.3 were observed (**Figure 1**). The ratio increased, as expected, when fast growing cells were suddenly starved for carbon (0.4), and especially nitrogen (0.8) (**Figure 1**). Deletion of *ATG1*, encoding a kinase essential for macroautophagy, abolished the response to starvation and ratios in fast growing and starved cells remained below 0.2 (**Figure 1**).

**Figure 1.**
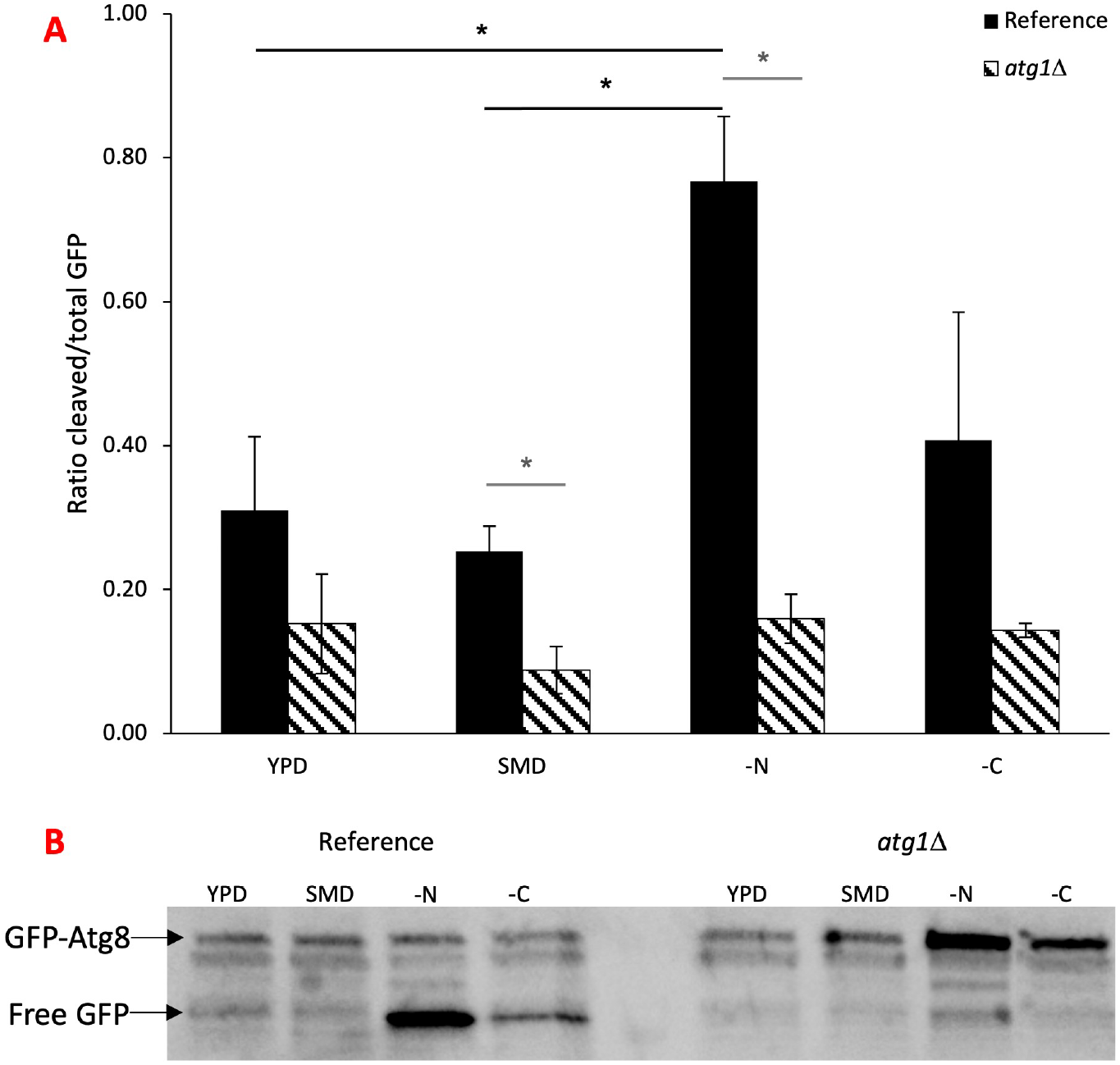
Cleavage of GFP-Atg8 in fast growing and starved yeast cultures. (**A**) Ratios of free GFP over total GFP (free GFP + GFP-Atg8) in growing cultures of the reference or *atg1*Δ strain on YPD or complete synthetic medium (SMD) or in cells starved for nitrogen (-N) or carbon (-C) for 4 hours. Data shown are the average of two duplicates, error bars indicate the range between replicates. *indicate significant differences (*p*-value < 0.05), between conditions for the reference strain (black lines), and between the reference and *atg1*Δ mutant strains (grey line). (**B**) Illustrative western blot showing GFP-Atg8 cleavage under different conditions for the reference and *atg1*Δ mutant strains used to determine the ratios shown in (A). Western blots used for quantification can be found in supplementary figure S1.

Subsequently, we monitored ratios in slowly to virtually non-dividing cells in retentostat cultures. In these continuously fed cultures, cells do not experience starvation. Their access to glucose is significantly restricted, but all other nutrients are present in excess, including the nitrogen source. The accumulation of initially dividing cells resulted in a further reduction in the glucose availability per cell and specific growth rates from 0.025 h^-1^ to 0.001 h^-1^, as observed previously (Vos et al. 2016). The ratios of free GFP over total GFP indicated that autophagy was active in aerobic retentostat cultures during the entire 14-day period (**Figure 2A, B**). Ratios were higher than in carbon-starved cells but lower than nitrogen-starved cells of the same strain (**Figure 1**). These results indicated that glucose-limitation induces autophagy already in dividing cells. This induction was not caused merely by the defined medium, as fast dividing cells on rich YPD and defined medium showed comparable low ratios of 0.3 (**Figure 1**).

**Figure 2.**
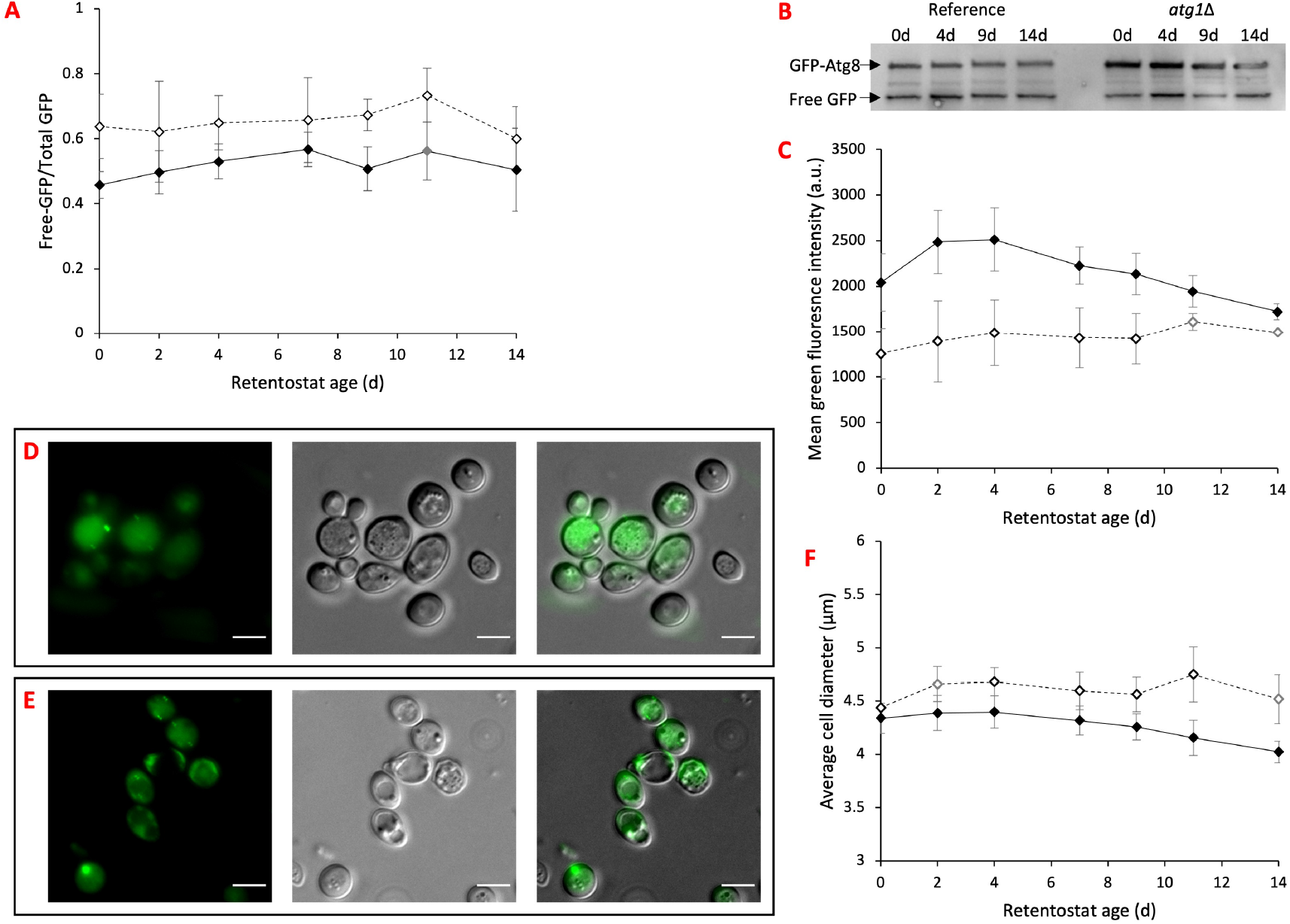
Autophagy in virtually non-dividing, non-starved cells of reference and *atg1*Δ mutant yeast strains. Autophagy was monitored by following cleavage, expression and localization of GFP-Atg8. Data shown are average values from four independent replicate retentostat cultures of the reference (open symbols) and *atg1*Δ mutant (closed symbols) strains, error bars indicate the standard deviation. (**A**) Ratios of free GFP over total GFP (free GFP + GFP-Atg8) in aging yeast cells. For age 11d, data of only triplicate *atg1*Δ cultures are shown, indicated in grey. Images of the western blots used for quantification can be found in supplementary figure S2. (**B**) Western blot image with GFP-Atg8 and free GFP bands illustrating the different processing between representative retentostat cultures of the reference and *atg1*Δ strains at ages 0, 4, 9 and 14d. (**C**) Green fluorescence intensities of cells as measured by flow cytometry corresponding to cellular GFP levels. Mean data for at least 20.000 cells per timepoint per replicate are shown. For age 11d and 14d data of only duplicate reference cultures are shown, indicated in grey. (**D**) Localization of GFP (green) in 14d old yeast reference retentostat cultures and (**E**) in 14d old *atg1*Δ mutant retentostat cultures: GFP, DIC and overlay. Scalebar corresponds to 5 μm. (**F**) Average size of cells in retentostat cultures. For age 2d and 14d data of only duplicate reference cultures are shown, indicated in grey.

Increasing retentostat culture age, i.e., while the division rates approached zero, did not impact GFP ratios (*p*-value > 0.5) (**Figure 2A, B**). To check whether the absence of a correlation between division rate and GFP ratios also held for autophagy fluxes, the fluorescence intensity of cells as a measure for total GFP levels were determined. These data also showed no change over time (**Figure 2C**, *p*-value > 0.3), suggesting that the autophagy flux remained constant under ECR.

### Absence of Atg1p does not abolish macroautophagy under extreme calorie restriction

To further explore the role and importance of autophagy in these aging retentostat cultures, we sought to abolish this recycling process. The highly conserved Atg1 plays a crucial role in autophagy initiation as well as phagophore assembly site organization and cycling of several other Atg proteins (Papinski et al. 2014; Cheong et al. 2008). Deletion of *ATG1* has been reported to result in defects in autophagy and shorter survival under starvation (Alvers et al. 2009). To monitor the effect of the *ATG1* deletion on autophagy activity in ECR cultures, free GFP/total GFP ratios were determined during the aging of retentostat cultures. Surprisingly, significant fractions of free GFP were still detected in the *atg1*Δ strain (**Figure 2A, B**). Ratios were however significantly lower than those observed for the reference strain (F(1,5):41.7, *p*-value = 0.001, **Figure 2A**). On the contrary, mean green fluorescence intensities per cell were higher in the *atg1*Δ mutant, indicating higher levels of total GFP (F(1,6):16.484, *p*-value = 0.007, **Figure 2C**). Morphological alterations were observed for the *atg1*Δ strain, including a significant decrease in cell size over time (F(1.8,5.4):19.1, *p*-value = 0.003). Beyond day 2, *atg1*Δ cells were on average smaller than those of the reference strain of the same age up to 13% (F(1,4):8.116, *p*-value = 0.046, **Figure 2F**). Furthermore, more GFP-foci were observed at what appears to be the border of vacuoles (**Figure 2E**). These observations were in stark contrast with those for batch cultures. Even though GFP levels increased when fast exponentially growing *atg1*Δ cells were subjected to nitrogen or carbon starvation, ratios of cleaved GFP over total GFP remained below 0.2 (**Figure 1**).

### Deletion of ATG1 results in increased maintenance energy consumption

Similar to the reference cultures, specific growth rates approached zero over the course of 14 days in *atg1*Δ retentostat cultures (**Figure 3A**). Absence of Atg1 did however strongly impact the apparent maintenance requirement (**Figure 3B**). Glucose maintenance requirements (*m_S_*) were 60% higher for *atg1*Δ cultures compared to reference cultures (10.4 ± 1.0 mg/g_cells_/h vs. 6.4 ± 0.7 mg/g_cells_/h, *p*-value = 0.003). Besides this increased maintenance requirement, autophagy-deficient cells also displayed a 2-fold higher death rate (k_d,_ 5.5 ± 1.2 · 10^-2^ d^-1^ vs 2.6 ± 0.5 · 10^-2^ d^-1^, *p*-value = 0.007) over the course of 14 days, illustrated by the stronger loss of viability (**Figure 3C**). A cause for this decrease in viability could be the increased maintenance requirement. Namely, if the glucose supply per cell would fall below the maintenance requirement, cells would eventually not have sufficient energy for homeostasis. Vos *et al.* (2016) developed a model to predict retentostat growth kinetics using the Pirt relation for the distribution of glucose between growth and maintenance (Pirt 1982). The scenario in which the glucose supply becomes insufficient to support homeostasis, would result in a negative growth-rate in this model. To test if this scenario could have occurred in our experiments, we applied the model to the used working volumes, flowrates and glucose concentrations for both strains. Based on this, our experimental set-up could actually have supplied cells with even higher m_S_ (≤12.5 ± 0.4 mg/g_cells_/h) with sufficient glucose. The glucose supply did thus not fall below the observed higher maintenance requirement.

**Figure 3.**
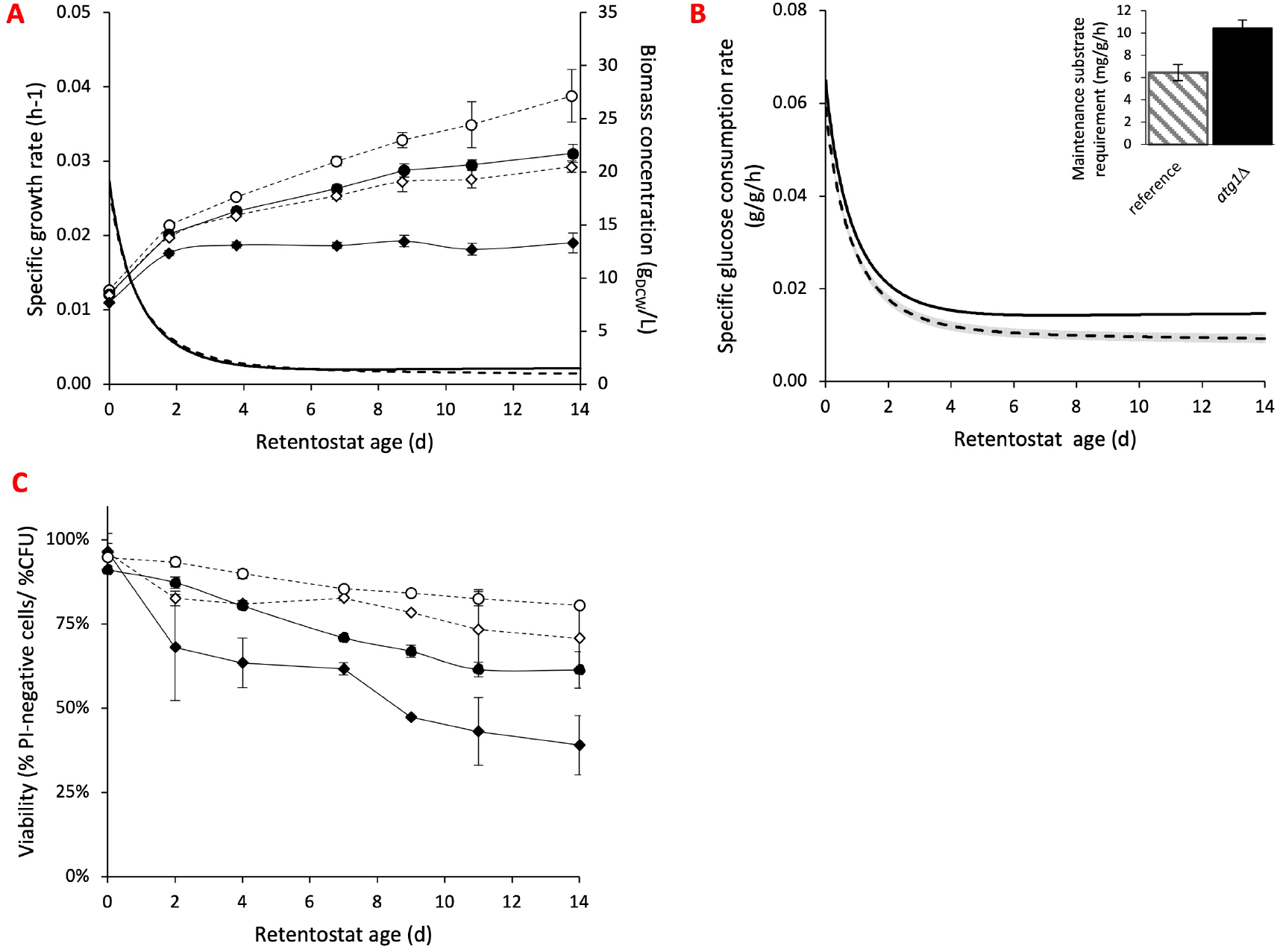
Effect of *ATG1* deletion on physiology of non-dividing yeast retentostat cultures. (**A**) Average biomass-specific growth rates of duplicate reference *ATG1* (dashed line) and *atg1*Δ mutant (continuous line) retentostat cultures (primary y-axis). Average total (circles) and viable (diamonds) biomass concentrations of *atg1*Δ (closed symbols) and reference (open symbols) quadruplicate cultures. Error bars indicate the standard deviation. (**B**) Specific glucose uptake rates of the *atg1*Δ mutant (continuous line) and reference strain (dashed line) approach the maintenance substrate requirements (*m_S_*, in set) while retentostat cultures age. Average values of two reference and four *atg1*Δ mutant independent replicate cultures are shown, error bars correspond to standard deviation. (**C**) Viability during the course of duplicate *atg1*Δ (closed symbols) and reference (open symbols) retentostat cultures as measured by CFU plating (diamonds) and PI staining (circles). Error bars correspond to standard deviation.

### Impact of *ATG1* deletion on mitochondrial function in respiring yeast cultures

Previous research has shown that under starvation, deletion of *ATG1* gives rise to a high fraction of respiratory deficient cells (Suzuki et al. 2011; Aris et al. 2013). Loss of respiration would present cells with a formidable challenge under retentostat conditions. The ECR results in a supplied amount of glucose that does not allow cells to generate sufficient ATP for maintenance by fermentation only. So, loss of respiratory capacity might explain both a higher average *m_S_* and reduced survival. Triphenyltetrazolium chloride (TTC) staining of colonies obtained in CFU assays revealed respiratory deficient cells in both reference and *atg1*Δ cultures (**Figure 4A**). Fractions of respiratory deficient cells were twice as high in the *atg1*Δ cultures, with on average 10.1% respiratory deficient cells in the *atg1*Δ mutant cultures versus 5.4% in the reference cultures. Over the culture duration, this difference was significant between the strains (F(1,5):9.6, *p-*value = 0.027). Prior to complete loss of respiratory capacity, respiratory efficiency might be lower due to reduced functionality of the electron transport chain and mitochondrial membrane leakage. Reduction in respiratory efficiency can affect the respiratory quotient (RQ), i.e., the amount of carbon dioxide produced per mole of oxygen consumed, as well as specific oxygen uptake rates. At the whole culture level, however, no differences in RQ were observed in the presence or absence of Atg1p. Specific oxygen uptake rates were 1.6 times higher for the *atg1*Δ strain compared to the reference strain in aged retentostat cultures (0.35 ± 0.02 vs 0.23 ± 0.01 mmol/g/h, *p*-value = 0.0009, **Figure 4B**). This increased oxygen consumption is mostly due to the maintained higher glucose catabolism. The amount of oxygen consumed per mole of glucose consumed was however also slightly, yet not significantly elevated, when autophagy was reduced (*p*-value 0.08, **Figure 4C**).

**Figure 4.**
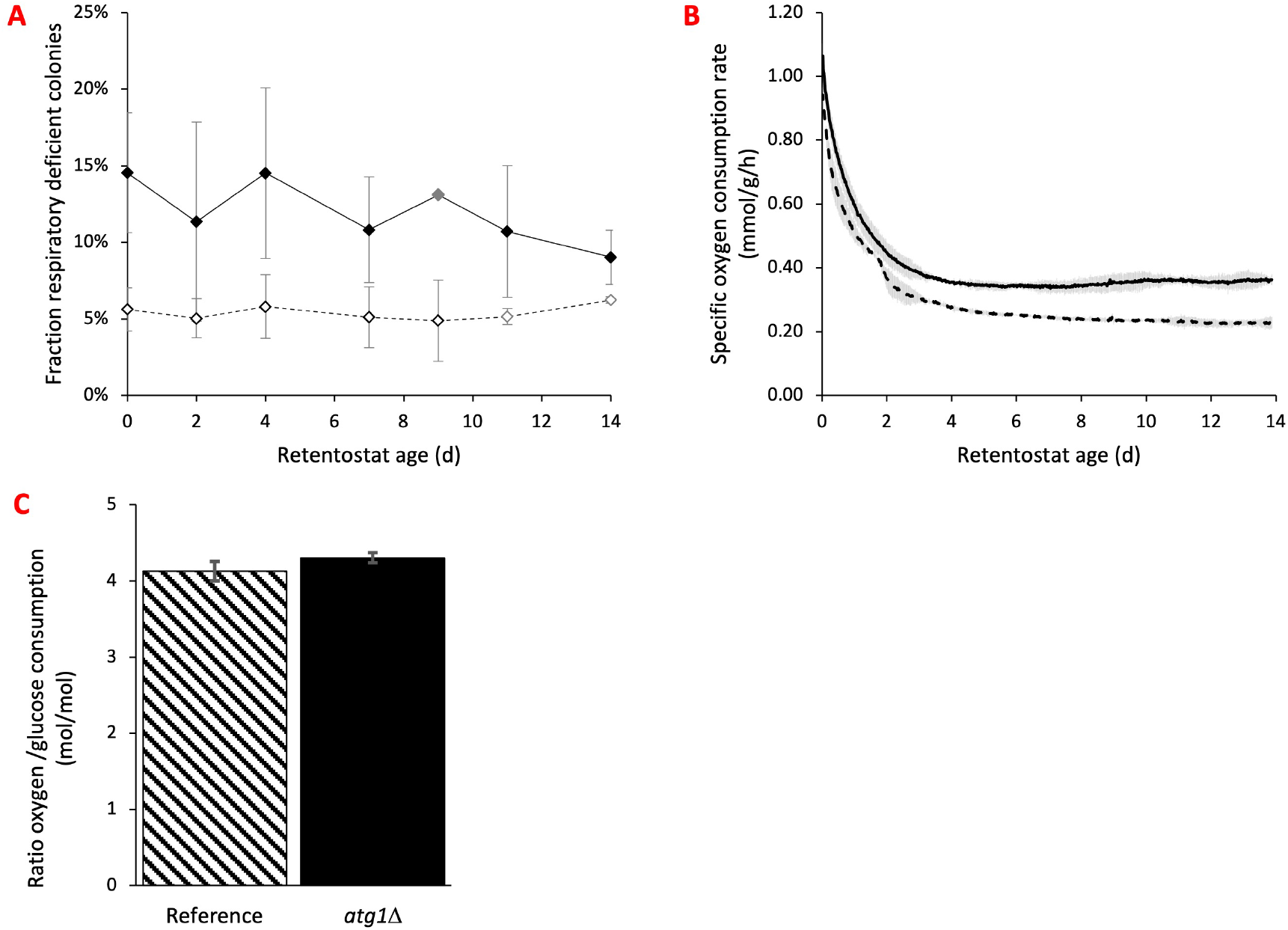
Absence of Atg1 affects respiratory capacity of virtually non-dividing, aging yeast cells. (**A**) Fraction of respiratory deficient cells as function of age in quadruplicate reference retentostat cultures (open symbols) and triplicate *atg1*Δ retentostat cultures (closed symbols) (age 9d was measured only in a single culture for the mutant strain, at age 11 and 14d values for duplicate refence cultures are shown, all indicated in grey). Respiratory capacity was determined using TTC staining on colonies from CFU determinations. (**B**) Average specific oxygen consumption rates of duplicate reference (dashed line) and *atg1*Δ (continuous line) retentostat cultures. Error bars correspond to standard deviation. (**C**) Ratio of the specific oxygen uptake rate over the specific glucose uptake rate for 14d old retentostat cultures. Values shown are averages of duplicate reference and quadruplicate mutant retentostat cultures, error bars represent standard deviation.

## Discussion

### Calorie restriction is a more potent inducer of autophagy than carbon starvation

Using continuously fed retentostats, yeast cultures can be brought into a maintenance-only state. In this state the consumption rate of glucose, the energy-substrate, just meets the maintenance energy requirement, leaving no additional energy for divisions and growth. This situation of ECR offers insights into which processes are active in such non-dividing, but non-starved cells. Such processes likely play an important role in actual cellular maintenance and aging. In this study we found that increasing ECR, monitored as the transition from slow to virtual zero growth, results in vacuolar accumulation of cleaved GFP-Atg8. These findings are in line with previous observations that carbon starvation, the most extreme case of calorie restriction, already results in higher autophagy activity (Iwama & Ohsumi 2019). Autophagy activity in starved cultures strongly dependents on the preceding growth conditions; lower initial glucose concentrations, often referred to as calorie restriction, stimulate autophagy and lead to yeast lifespan extension (Aris et al. 2013). The physiological state of cells in retentostat cultures differs in important aspects from their starved counterparts (Bisschops et al. 2017; Boender, Almering, et al. 2011). This difference in physiological state is also reflected in the differences in GFP-Atg8 cleavage between abruptly carbon-starved cells and cells in retentostat cultures, where glucose uptake and growth rate decrease gradually (**Figure 1 and 2A**). Intriguingly the degree of energy-limitation did not affect the ratio nor total levels of cleaved GFP-Atg8 in the range tested here (**Figure 2A and 2B**). This absence of correlation between energy-limitation and autophagy flux is not because maximal delivery and cleavage of GFP-Atg8 were reached. Higher cleaved:total ratios of GFP-Atg8 were observed, when the same strain was subjected to nitrogen-starvation, one of the most potent autophagy inducers in yeast (Delorme-Axford & Klionsky 2018) (**Figure 1**).

### Energy-saving by autophagy in aging yeast cells is only partially due to improved mitochondrial homeostasis

A unique aspect of retentostat cultures is that they allow very accurate quantitative estimation of the growth-rate independent maintenance energy requirement. Deletion of *ATG1* led to a 60% increase in maintenance energy requirement (**Figure 3B**). As the maintenance energy requirements were based on glucose consumption, rather than actual ATP conversion rates, this increase in glucose consumption for maintenance might be explained for by a loss or reduction of respiratory efficiency. Loss of respiration results in an 8-fold lower ATP yield per glucose catabolized. The observed difference between (average) 5% respiratory deficient cells in reference cultures and 10% in the *atg1*Δ cultures would result in at most a 30% increase of the average maintenance substrate consumption. Autophagy thus allows yeast cells to economize their energy substrate, not only by improved maintenance of mitochondrial functionality, but also in other yet unknown ways. A likely explanation may lie in the recycling role of autophagy, which may reduce energy expenditure on especially amino acid synthesis.

Besides recycling of macromolecules, autophagy also plays a vital role in healthy aging through turnover of damaged macromolecules and organelles (Sampaio-Marques et al. 2014). The role of autophagy should therefore be more prominent with increasing age of the culture. This is indeed what we observed. Although a slightly elevated fraction of dead cells was already observed while cells were slowly dividing (*p*-value = 0.01 at t = 0) in *atg1*Δ cultures, the difference in survival became more apparent when division ceased and the cells in the cultures aged (**Figure 3C**). Concomitantly with chronological aging, the availability of energy (glucose) decreased (**Figure 3B**). If the cellular maintenance requirement is growth-rate and age independent, this will eventually result in an equilibrium between the specific growth-rate and death rate: Each cell in the reactor that dies and hence stops consuming energy substrate, leaves a bit more substrate to be taken up by the remaining cells which can divide as the net uptake supersedes the maintenance energy requirement, albeit only minimally. For the reference strain this holds in the timespan assessed here, the specific growth rate approached the specific death rate, but did not fall below (*μ_end_* = 0.0014 ± 0.0002 h^-1^ vs k_d_ = 0.0011 ± 0.0002 h^-1^). This can also be seen by the slight increase in viable biomass during the course of the culture (**Figure 3A**). For the *atg1*Δ cultures this did not hold and the specific growth rate decreased to below the specific death rate (*μ_end_* = 0.0022 ± 0.0004 h^-1^ vs k_d_ = 0.0023 ± 0.0005 h^-1^), resulting in a slight decrease in viable biomass concentration (**Figure 3A**). This implies that besides an already higher initial maintenance substrate requirement, this requirement increases with age. In contrast, the fraction of respiratory deficient cells did not increase with age. This supports our notion that the increased glucose needs for maintenance due to impairment of autophagy, not only result from catabolic challenges.

### Loss of respiratory capacity is not the only driver of loss of viability

Previous studies showed that *atg1*Δ and other autophagy mutants survive shorter in stationary phase cultures. Here we observed a doubling of the death rate (*kd)*, which shows that even in the presence of an extracellular energy source and a surplus of nitrogen, impairment of autophagy results in accelerated cell death. Although the fraction of viable cells that lost respiratory capacity showed no correlation with culture age for both the reference and *atg1*Δ strains, the fraction of respiratory deficient cells was also twice as high in the *atg1*Δ strain retentostat cultures. Based on these two observations, one could hypothesize that loss of respiratory capacity dictates loss of viability. In this case, the chronological lifespan of respiratory deficient cells would be independent of Atg1 expression. Combining the retentostat data presented here, the average CLS of respiratory deficient cells, i.e., the time of survival after respiratory capacity has been lost, can be calculated. The respiratory deficient cells (*C_x,resp def_*) in this case would be the cells to lose viability, i.e., the change in dead cell concentration (d*C_x,dead_*) (Eq.1). The equation to estimate the death rate (*k_d_*) as defined by Vos *et al*. (2016) (Eq. 2) can then be combined with equation 1 to calculate the CLS of respiratory deficient cells (Eq. 3).

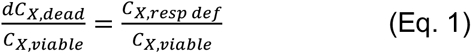

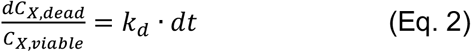

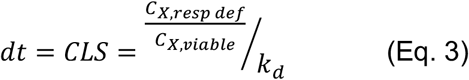

Based on these equations the estimation of the average CLS of respiratory deficient cells of the reference strain and of the *atg1*Δ mutant strain is indeed similar with resp. 49h and 44h. Compared to previous data from retentostat and other cultures, a CLS of 2 days is however unlikely short, even for respiratory deficient cells. For example, under anaerobic conditions (no oxygen for respiration), the same reference strain displayed a CLS of 4 days in stationary phase cultures and even longer in starved retentostat cultures (Bisschops et al. 2015; Boender, Almering, et al. 2011). Cells in active retentostats however have been preconditioned, and contain high levels of intracellular reserves and still consume a limited supply of glucose (Vos et al. 2016), factors that would rather elongate than shorten CLS (Boender, Almering, et al. 2011). These observations thus imply that the fraction of respiratory deficient cells is not equal to the change in dead biomass (Eq. 1) and, hence, that loss of respiratory capacity is not the main, direct cause of loss of viability in retentostat cultures. Furthermore, the CLS of respiratory deficient cells calculated using Eq. 3 was almost equal for both strains. The *atg1*Δ mutant, however, has a 60% higher maintenance requirement, which can only be partially attributed to the higher fraction of respiratory deficient cells. An increased maintenance requirement likely results in a reduced CLS, because energy reserves will be depleted faster, and cells experience a stronger and earlier energetic crisis to sustain homeostasis. This thereby also argues against the hypothesis that loss of respiratory capacity per se is the major cause of death in aging retentostat cultures.

### Maintained autophagy in the absence of Atg1 under ECR and chronological aging conditions

The kinase Atg1 plays an essential role in macroautophagy, which relies on Atg8 for phagophore development into autophagosomes and expansion thereof. In surplus of all other nutrients, autophagy activation was previously only observed after glucose depletion, when cells started and continued to respire ethanol (Iwama & Ohsumi 2019). Here we show that full respiration of glucose also results in activation of autophagy. The observed cleavage of GFP-Atg8 in yeast retentostat cultures is at least in part due to macroautophagy. However, in the absence of Atg1, degradation of Atg8 still occurred, resulting in free GFP. Recently, Iwama and coworkers also observed cleavage of GFP-Atg8 in post-diauxic shift cultures, i.e., fully respiring cells, of several *ATG* mutants, including *atg1* and *atg2* (Iwama & Ohsumi 2019). They showed that under these conditions, microautophagy is also active and results in the breakdown of Atg8. Other authors previously demonstrated the activation of microautophagy during and after the diauxic shift in yeast batch cultures (Oku et al. 2017). In macroautophagy, an autophagosome forms around the cargo to be degraded and this double-membrane structure subsequently fuses with the vacuole. Microautophagy on the other hand occurs through invaginations of the vacuolar membrane directly (Oku et al. 2017). Retentostat cultivation captures yeast cells for prolonged periods of time in a state that shares important metabolic features with post-diauxic shift cultures, like fully respiratory and with (close to) zero glucose repression (Vos et al. 2016; Boender et al. 2009). Hence the cleavage of GFP-Atg8 in *atg1*Δ retentostat cultures could be the result of microautophagy. The microscopic observation of fluorescent foci at the vacuolar membrane (**Figure 2D**) support this. Microautophagy is to date lesser studied, but is also a conserved mechanism shared between yeast and human cells (Mejlvang et al. 2018). In human cells, microautophagy has also been shown to be involved in degradation of the Atg8-like proteins LC3B and GABARAPL2 (Mejlvang et al. 2018).

### Concluding remarks

Here we showed that under prolonged ECR, autophagy is active and contributes to cellular survival. Alterations in autophagy, such as deletion of crucial components of the macroautophagy machinery strongly affect cellular energy-substrate requirements. These findings have important implications for the most used system to study chronological aging in yeast: stationary phase cultures. In these nutrient-depleted cultures, the shortened survival of autophagy mutants may not only be the consequence of enhanced aging, but also of more rapid depletion of limited intracellular reserves. Combined with observations in the present study and previous work that impaired autophagy increases petite formation and that autophagy plays a role in lipid metabolism, cells with inefficient autophagy machinery may rapidly run into an energetic crisis, even before (or strengthened by) cellular aging. Retentostats in this light provide an additional tool to study chronological aging at the cellular level. Also in these cultures, cell division comes to a virtual standstill and, due to the cell retention, cell death can be monitored over time. The continuous supply of glucose (in restriction) and all other essential nutrients (in excess), however results in a different physiological state and can provide deeper insight in the links between cell death, aging and energetics.

Further investigations are needed to elucidate which other roles, besides mitochondrial homeostasis, contribute to the positive effects of autophagy on chronological lifespan. Also, in addressing whether (stronger) activation of Atg1-independent autophagy is caused by reduction of macroautophagy or other processes and which genetic factors are involved, retentostats can play a pivotal role.

### Experimental procedures

#### Yeast Strains and media

The prototrophic *S. cerevisiae* strain IMX585 (*MATa can1* Δ::*cas9*-*natNT2 URA3 TRP1 LEU2 HIS3*) (Mans et al. 2015) harboring the *cas9* gene and derived from the model strain CEN.PK113-7D (Nijkamp et al. 2012) was used for the construction of reference strain IMI442 (*MAT*a *can1*Δ::*cas9*-*natNT2 URA3 TRP1 LEU2 HIS3 X-2*::*gfp-ATG8*) and the *atg1*Δ mutant IMK907 (*MATa* URA3 *can1*Δ::*cas9*-*natNT2 TRP1 LEU2 HIS3 X-2*::*gfp*-*ATG8 atg1*Δ). The *gfp*-*ATG8* expression cassette was amplified from plasmid pRS416-*GFP-ATG8* (a gift from Daniel Klionsky, addgene #49425) (Guan et al. 2001) with 60bp flanks homologous to the integration site. The integration site on chromosome X, X-2, was chosen based on (Mikkelsen et al. 2012). For efficient homologous recombination guided integration, a double-strand break in the X-2 region was induced by a gRNA-Cas9 complex following the method described by (Mans et al. 2015). The X-2 targeting gRNA was expressed from a plasmid carrying the spacer CTACTCTCTTCCTAGTCGCC constructed from pMEL13 (a gift from Jean-Marc Daran, addgene #107919). For marker-free deletion of *ATG1*, two gRNAs targeting the *ATG1* coding sequence, with spacers CAACAGTATTTTCAAAATCA & CGTTAGTATTTTGGAATCTC, were expressed from a plasmid based on pROS13 (a gift from Jean-Marc Daran, addgene #107927). Target sites and repair fragments were selected and designed based on the yeastriction webtool (Mans et al. 2015). Yeast transformations were done according to the lithium-acetate, heat-shock method (Gietz & Schiestl 2007) and genetic modifications were confirmed by diagnostic PCR. Strains were grown until early stationary phase in YPD medium (10 g/L yeast extract, 20 g/L peptone and 20 g/L dextrose) and stored with 20% glycerol at −80°C as cryostocks.

Yeast cells were cultured in defined medium as described by (Verduyn et al. 1992) containing 7.5 to 20 g/L glucose as sole carbon-source (see below). For bioreactor cultivations, the defined medium was supplemented with 0.25 g/L Pluronic 6100PE (BASF, Germany) to prevent foaming. In the medium containing 20 g/L glucose the concentrations of trace elements and vitamins were increased, 1.5 and 2.0-fold respectively as described by (Vos et al. 2016).

#### Chemostat and Retentostat cultivations

Aerobic chemostat and retentostat cultivations were essentially performed as described earlier (Vos et al. 2016). 2-liter bioreactors (Applikon, The Netherlands) were operated at a working volume of 1.4 L and a dilution rate of 0.025 h^-1^. Temperature was kept constant at 30°C and pH was controlled at 5 by automatic addition of 10% ammonium hydroxide. Cultures were sparged with dried air at 0.5 vvm and continuously stirred at 800 RPM. Medium was supplied to the bioreactor from a stirred (500 RPM) mixing vessel (working volume 1.2 L, Applikon, The Netherlands). The mixing vessel was fed from medium reservoirs containing defined medium containing 20 g/L or 7.5 g/L glucose as carbon-source. Cultures were considered in steady state when, after at least 6 volume changes, biomass concentrations and CO2 production rates differed less than 5% over 2 volume changes.

Retentostat cultures were started from steady-state chemostat cultures by redirecting the effluent through a port equipped with a polypropylene filter of pore-size 0.22 μm (Applisense, Applikon, The Netherlands) prepared as described earlier (Vos et al. 2016). Simultaneously the feed to the mixing vessel was changed from 20 g/L glucose medium to 7.5 g/L glucose medium and pH was controlled at 5 by the addition of 2M KOH. Full retention of biomass was verified by plating of filter effluent on YPD at least at 2 different timepoints per cultivation. Biological quadruplicates were run for each strain.

#### Starvation experiments

To induce autophagy for reference data, yeast cells were grown in round-bottom shake flasks, working volume 100 mL, at 30°C and 200 RPM in Innova 44 incubators (New Brunswick, USA). Cultures were inoculated from cryostocks. After 8 hours cells were transferred to fresh media to obtain exponentially growing cultures overnight. Exponentially growing cells were centrifuged and transferred to tubes containing 10 mL of YPD, defined medium with 20 g/L glucose, defined medium lacking glucose (carbon starvation) or defined medium lacking ammonium sulfate (nitrogen starvation) at a starting OD_660_ of 10. After 4 hours incubation at 30°C, samples corresponding to approximately 25 mg of dry cell weight were taken for western blot analysis.

#### Biomass, substrate, extracellular metabolite and gas concentration analyses

Biomass concentrations as concentrations of dry cell weight were determined as described previously (Vos et al. 2016). In short, exactly 10 mL of an appropriate dilution of culture broth was filtered through a pre-dried polyether sulfone filter (pore size 0.45 μm, Pall Corp., UK). Filters were washed with demineralized water and dried in a microwave oven (360 W, 20 min). The increase in mass of the filter corresponds to the dry cell weight.

Cell concentrations and volumes were determined using a Z2 Coulter Counter (50 μm orifice, Beckman Coulter, The Netherlands).

For substrate and extracellular metabolite concentrations determination samples were rapidly cooled and centrifuged (5 min, > 5000 g at 0°C) prior to HPLC analysis as described previously (Bisschops et al. 2017).

In- and outgoing gas of bioreactors was dried prior to analysis of O_2_ and CO_2_ fractions using a Rosemount NGA 2000 analyzer (Emerson, USA). Off-gas was cooled to 2°C before leaving the bioreactor to prevent evaporation. Total oxygen consumption rates (R_O2_) were calculated by multiplying the difference in oxygen fraction between the off-gas and the in-gas with the flow rate (in mol/h). For specific oxygen consumption rates (q_O2_), total oxygen consumption rates (R_O2_) are divided by the amount of viable biomass in the reactor.

#### Viability measurements and TTC staining

Viability of cells was assessed by propidium iodide (PI) staining and colony forming unit (CFU) counting. PI stains cells with a permeable membrane, which are considered dead (Carmona-Gutierrez et al. 2018). For the PI assay, cells were diluted to 1.0 × 10^7^ cells/mL suspensions in Isoton II Diluent (Beckman Coulter, The Netherlands). Cells were incubated in the presence or absence of 0.02 mM PI in the dark at room temperature for 20 minutes. At least 10.000 cells were analysed for red fluorescence (exc. 488nm, FL-3, 670 nm LP filter) on a BD-Accuri C6 flowcytometer (Becton Dickinson, USA). Viability is shown as the fraction of PI-negative cells.

Cells for CFU assays were sampled from bioreactors directly into 10 mM Na-Hepes pH 7.2 buffer with 20 g/L glucose to prevent starvation (Boender, van Maris, et al. 2011). Cell concentrations in the suspensions were determined as described above. Cells were diluted in 0.1% peptone and 100 uL aliquots containing 200-300 and 20-30 cells were plated on YPD agar plates in triplicate. Colonies were scored after 2-3 days incubation at 30°C. Viability is depicted as the number of CFU divided by the number of plated cells.

Respiratory deficient colonies were identified using the 2,3,5-triphenyltetrazolium chloride (TTC) overlay technique (Ogur et al. 1957). Respiratory competent cells reduce TTC and turn red. YPD plates from CFU analysis were covered with defined medium agar containing 10 mM TTC. After solidification of the agar, plates were incubated at room temperature at least overnight. White colonies were scored as respiratory deficient and shown as the fraction of the total number of colonies.

#### Western blot analysis

For GFP and GFP-Atg8 analyses by western blot, 25 mg of cells were sampled from bioreactors and immediately cooled to 0°C. Cells were washed twice with ice-cold demineralized water (5 min, > 5000g at 0°C) and pellets were stored at −80°C. Cell free extract preparation was essentially performed as described by Chen and Petranovic (Chen & Petranovic 2015). Cell pellets were resuspended in 200 μL ice-cold lysis buffer (50 mM HEPES pH 7.5, 150 mM NaCl, 2.5 mM EDTA, 1% v/v Triton X-100, Complete Mini Protease Inhibitors (Roche, Switzerland)) in safe-lock tubes (Eppendorf, Germany). 100 μL of acid-washed glass beads (425 - 600 micron, Sigma, USA) were added and the suspension was homogenized using a FastPrep-24 (MP Biomedicals, France) in three rounds of 45s shaking at 6.5 m/s and 5 min cooling on ice. Supernatants were cleared in three centrifugation steps (20.000g 10 min at 0°C). Protein concentrations were determined using the Quickstart Bradford Protein Assay (Biorad, USA). 25 μg of total protein was separated on 4-15% Mini-Protean TGX Stain-Free Protein gels (Biorad, USA) at 250 V for 20 min. Gels were activated to check the quality of the protein extracts using the stain free protocol on a ChemiDoc MP Imager (Biorad, USA). Proteins were transferred to PVDF membranes using a Turboblotter system (Biorad USA). Successful transfer of proteins was verified using the ChemiDoc MP Imager. Membranes were blocked for 1 hour in TBS + 1% Casein (Blocking Buffer, Biorad, USA) and washed once with PBSt (Phosphate saline buffer pH 7.2 with 0.05% Tween20). Washed membranes were incubated overnight, shaken at 4°C, in PBSt with anti-GFP antibody (1:1000, from mouse, Roche, Switzerland). Membranes were washed in PBSt and incubated with the secondary anti-mouse HRP antibody (1:2000, from goat, Dako Agilent, USA) for 3 hours at 4°C. Subsequently, membranes were incubated with anti-GAPDH Alexa Fluor 647 antibody for 1 h at room temperature. Enhanced chemiluminescence signals were detected using the ECL Prime reagent (GE Healthcare, USA) and the ChemiDoc XRS imaging system (BioRad, USA). Binding was detected as chemiluminescence signals using ECL Prime reagent (GE Healthcare, USA) or fluorescence signals using a ChemiDoc MP Imager (BioRad, USA). Images were analyzed and bands were quantified using Image Lab 6.1 software (Biorad, USA).

#### Fluorescence detection

GFP fluorescence was detected using microscopy and flow cytometry. For flow cytometry, cells were diluted to 1.0 × 10^7^ cells/mL suspensions in Isoton II Diluent (Beckman Coulter, The Netherlands). At least 10.000 cells were analyzed for green fluorescence (exc. 488nm, FL-1, 530/30 nm BP filter) on a BD-Accuri C6 flow cytometer (Becton Dickinson, USA).

For fluorescence microscopy, cells were imaged using a Zeiss D1 Imager (Zeiss, Germany) equipped with a Plan-Apochromat 63x/1.40 Oil DIC M27 objective (Zeiss, Germany). GFP was excited using a HAL100 fluorescent lamp and filter set 10 (Ex 450–490 nm/Em 515–565 nm) (Zeiss, Germany). DIC and green fluorescence images were taken with an AxioCam HRm camera (Zeiss, Jena, Germany). Images were analysed using Fiji version 2.0.0-rc-69/1.52p (NIH, USA).

#### Regression and statistical analyses

Estimation of specific growth, glucose consumption and death rates as well as substrate maintenance requirements in individual aerobic retentostat cultures was performed using the least-squares regression analysis described by (Vos et al. 2016). Using determined biomass, PI-staining based viable biomass and glucose concentrations as input parameters along with measured cultivation parameters (reactor and mixing vessel volumes, flowrates) and a previously estimated *Y_sx_^max^* of 0.5 g/g.

To verify whether observed increased maintenance requirements could have resulted in starvation, i.e., specific glucose consumption rates below the maintenance requirement, the prediction of retentostat growth kinetics as performed by Vos *et al.* (2016) for the parental strain was repeated using a higher maintenance requirement as input. Outcomes of the simulations were checked for negative specific growth-rates while using the variables as used during the cultivation (reactor and mixing vessel volumes, glucose concentrations, flowrates and *Y_sx_^max^* of 0.5 g/g). All simulations and models were run using MATLAB version R2020a (The Mathworks, USA).

Pairwise comparisons were tested using a standard two-sided Student’s *t*-test assuming homoscedastic variance including data from all biological replicates. For time-course comparisons per strain, repeated measures ANOVA was performed, for time-course comparisons between strains, mixed ANOVA was performed including the time-points with 3 or 4 replicate measurements. ANOVA was performed using the SPSS software (v28.0.1.0, IBM Corp) and significant results are reported as F-value (degrees of freedom (df), error in degrees of freedom): *p*-value). Main effects were compared using a Bonferroni correction. For time-course analyses the values resulting after Greenhouse-Geisser correction are reported. A *p*-value of 0.050 was used as an upper threshold to reject the null hypothesis that no difference exists.

## Acknowledgements

The authors would like to thank Pascale Daran-Lapujade and Jack Pronk for valuable feedback during the work and Erik de Hulster and Marijke Luttik for technical assistance in setting up cultivation and analytical methods at TU Delft.

The work performed at Chalmers University by XC, AvA, DP and MB was funded by the Novo Nordisk Foundation center for Biosustainability with grant 21210022. The work in Delft by AvA and MB was funded by the Dutch Research Council (NWO) by grant 016.Veni.181.056 and equipment was sponsored by gifts to the Delft University Fund.

## Author’s contributions

MB, XC, AvA designed and performed experiments and data analyses. MB and DP designed the study and acquired funding. MB wrote the manuscript. All authors contributed to data interpretation and manuscript revision.

## Data availability

All raw data as well as used strains are available upon request.

## Supplementary materials

<u>Supplementary Figure S1.</u> Western blots used to determine the ratio between free and total GFP in exponentially growing and starved cultures shown in Figure 1.

<u>Supplementary Figure S2.</u> Western blots used to determine the ratio between free and total GFP in retentostat cultures shown in Figure 2.

**Supplementary figure S1.**
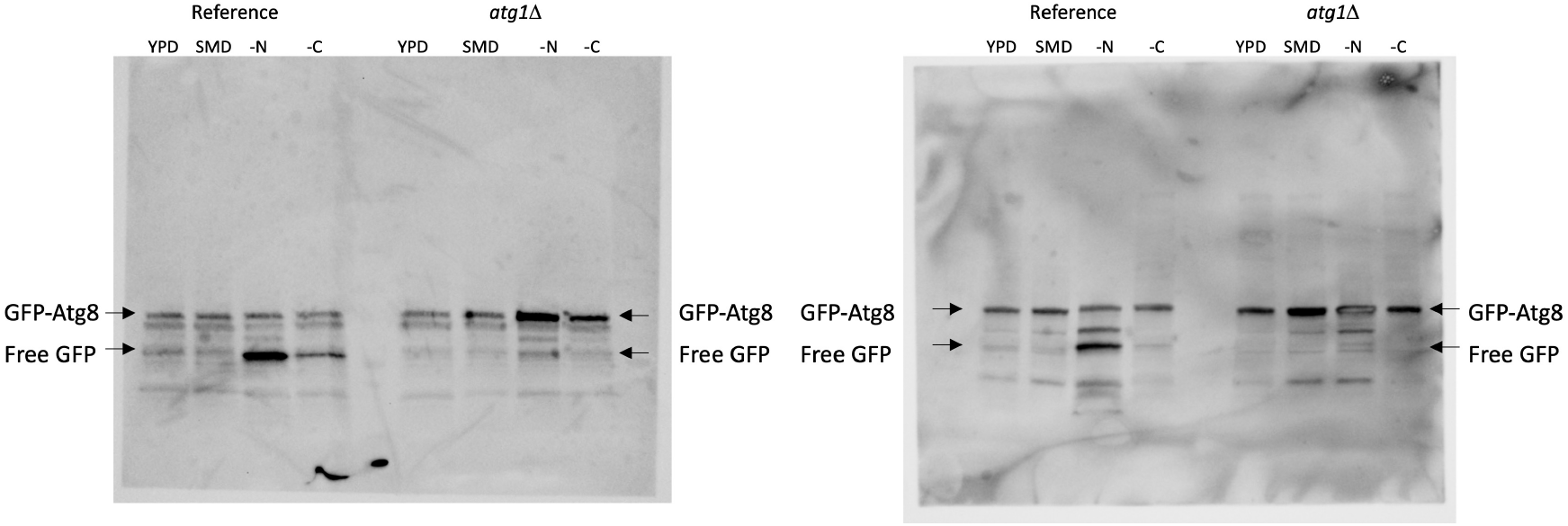
Western blot images of the chemiluminescent signals used for figure 1. Cultures of the reference strain and the *atg1*Δ strain are shown. YPD: exponential growing cells on rich YPD medium, SMD: Exponential growing cells on chemically defined medium with glucose, -N: Cells starved for 4 hours in defined medium with glucose lacking nitrogen, -C: Cells starved for 4 hours in defined medium without glucose.

**Supplementary figure S2.**
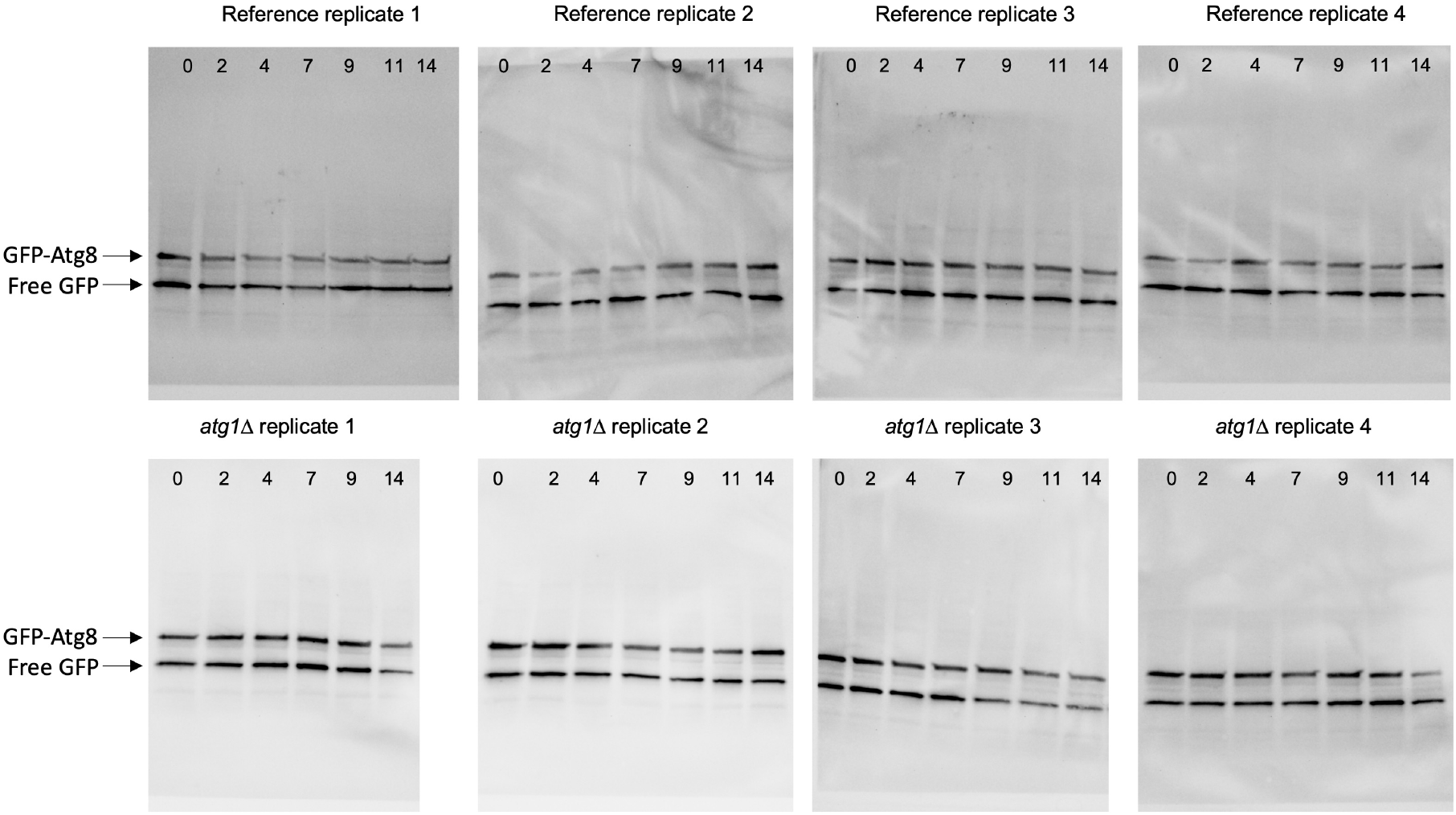
Raw Western blot images of the chemiluminescent signal per retentostat culture used for figure 2. Top row contains the images for the reference strain cultures, bottom row for the *atg1*Δ strain cultures. Numbers indicate the retentostat age in days.

